# Fetal alleles predisposing to metabolically favourable adiposity are associated with higher birth weight

**DOI:** 10.1101/2020.09.17.302208

**Authors:** WD Thompson, RN Beaumont, A Kuang, NM Warrington, Y Ji, J Tyrrell, AR Wood, D Scholtens, BA Knight, DM Evans, WL Lowe, G Santorelli, R Azad, D Mason, AT Hattersley, TM Frayling, H Yaghootkar, MC Borges, DA Lawlor, RM Freathy

## Abstract

**Background:** Higher birth weight is associated with higher adult body mass index (BMI). If genetic variants can be identified with alleles that predispose to both greater fetal growth and to greater adult adiposity, such shared genetic effects might indicate biological processes important in the early patterning of adiposity. However, variants identified in genome-wide association studies of adult BMI have overall been only weakly associated with birth weight. Genetic variants have recently been identified where one allele is associated with higher adult body fat percentage, but lower risk of metabolic disease, likely due to a favourable body fat distribution. The effect of these adult metabolically favourable adiposity alleles on an individual’s own birth weight is unknown.

**Aim:** We aimed to test the effect on birth weight of a fetal genetic predisposition to higher metabolically favourable adult adiposity and to compare this with the effects of a fetal genetic predisposition to higher adult BMI. We also aimed to examine the effects of a genetic predisposition to higher metabolically favourable adult adiposity or BMI on other birth anthropometric traits (length, ponderal index, head circumference and skinfold thickness) and on cord-blood insulin, leptin and adiponectin.

**Methods:** We used published GWAS data from up to 406,063 individuals to estimate the fetal effects on birth weight of alleles that are robustly associated with higher metabolically favourable adult adiposity or BMI. We additionally used 9,350 mother-child pairs from four cohorts to test the effects of the same alleles on other birth anthropometric traits and cord-blood markers. In all analyses, we adjusted for potential confounding due to the maternal genotype. We used inverse-variance weighted meta-analyses to combine summary data across SNPs.

**Results:** Fetal genetic predisposition to higher metabolically favourable adult adiposity was associated with higher birth weight (10 grams (95% CI: 7 to 13) higher mean birth weight per 1 SD pooled “genetic score”). Fetal genetic predisposition to higher adult BMI was also associated with higher birth weight, but with a smaller magnitude of effect (4 grams (95% CI: 0 to 8) higher mean birth weight per 1 SD pooled “genetic score”) and with higher heterogeneity across SNPs. Effects on other birth anthropometric outcomes were consistent with the effect on birth weight but with wider confidence intervals. There was no strong evidence for an effect on cord-blood markers.

**Conclusions:** Some genetic variants previously linked to adult adiposity influence birth weight. Alleles that predispose to higher metabolically favourable adult adiposity collectively have a stronger effect on birth weight than those predisposing to higher BMI. This suggests that the early accumulation of a metabolically favourable fat distribution might underlie part of the observed association between higher birth weight and higher adult BMI. Larger samples are needed to clarify the effects on other birth anthropometric measures and cord-blood markers.

## Introduction

High birth weight (> 4kg), compared to average birth weight, is associated with an increased risk of higher adult BMI[1]. Moreover, across the distribution of birth weight, higher birth weight is associated with higher BMI between the ages of 18 and 20 years[2]. The mechanisms underlying the association between higher birth weight and higher adult BMI are not fully understood, but one possible mechanism could be the inheritance of genetic variants that influence both fetal and postnatal body mass accumulation (particularly of body fat).

A previous large study of twins did not support a strong overall contribution of genetics to the correlation between birth weight and BMI, as it found that the intra pair association of birth weight and BMI was similar in both monozygotic and dizygotic twins[3]. In the most recent GWAS of birth weight[4], two of the 190 loci identified were also associated with adult BMI. However, it was uncertain whether the birth weight effects at the two loci were direct fetal effects, or maternal genetic effects influencing birth weight indirectly via the intrauterine environment.

Previous studies of European ancestry participants suggested that genetic variants that influence adult BMI have no association with birth weight[5-7]. Recently, however, using a much larger population sample than previous studies, a positive genetic correlation between birth weight and adult BMI was detected[4]. Subsequent studies have shown that a BMI polygenic risk score, derived from genetic variants reaching genome wide significance in a large genome wide study of BMI[8], is predictive of birth weight (the top 10% of BMI polygenic risk having a 60g higher birth weight than the bottom 10%)[9], suggesting that some of the fetal genetic predisposition to higher adult adiposity may influence birth weight.

Several genetic variants have been identified where one allele is associated with higher adult adiposity, but lower metabolic risk (i.e. lower risk of type II diabetes and cardiovascular disease), so called “metabolically favourable adiposity” alleles[10]. Metabolically favourable adiposity alleles may have this effect because they are associated with increased adiposity in the more metabolically stable subcutaneous adipose tissue and decreased fat deposition in the liver[10,11].

The primary aim of this study was to test the hypothesis that fetal genetic predisposition to adult metabolically favourable or general adiposity (the latter indexed by BMI) influences birth weight. To the best of our knowledge this is the largest sample size used (N = 406,063) to test whether fetal genetic predisposition to higher adult BMI affects birth weight, and the first to test the effect of fetal genetic predisposition for higher metabolically favourable adult adiposity on birth weight. A secondary aim of this study was to test the effects of the same fetal genetic predisposition to higher metabolically favourable adult adiposity or BMI on other perinatal anthropometric traits (length, ponderal index, head circumference and skinfold thickness) and cord-blood markers (insulin, c-peptide, leptin and adiponectin). This secondary aim was exploratory, given the relatively small sample sizes that we have available (N = 9,350).

## Methods

Our analyses of genetic effects on birth weight used publicly available Early Growth Genetics consortium (EGG)+UK Biobank summary genome wide association study (GWAS) data. We performed analyses of other birth outcomes using individual level data from four birth cohorts, each analysed separately, with results pooled using fixed effect meta-analysis. We limited all analyses to participants of European ancestry, as genetic variants related to metabolically favourable adiposity and BMI were discovered in GWAS of European ancestry individuals. We adjusted all analyses for maternal genotype effects to avoid confounding, since maternal and offspring genotypes are correlated and maternal genetic variants related to metabolically favourable adiposity or BMI are known to affect offspring birth weight[4,12,13].

### Data Sources

#### EGG+UK Biobank

The EGG consortium component of the GWAS data for birth weight used in this study includes a meta-analysis of 35 studies of fetal genotype with birth weight (N = 80,745, white European ancestry) as well as 12 studies of maternal genotype with offspring birth weight (N = 19,861, white European ancestry), with some of the fetal and maternal studies overlapping[4]. These studies were further meta-analysed with summary data on white European participants’ from the UK Biobank, which made up approximately 90% of the meta-analyses (with N = 217,397 contributing to fetal genotype analyses and N = 190,406 contributing to maternal genotype analyses). Between 2006 and 2010, UK Biobank participants were recruited from the NHS patient registers and contacted if they lived in close proximity to one of 22 assessment centres in England, Scotland and Wales. Detailed medical data was collected on 502,655 participants (5.5% response), aged between 40 and 69 at recruitment[14]. All participants provided written informed consent, including for their collected data to be used by international scientists. UK Biobank has approval from the North West Multi-centre Research Ethics Committee (MREC), which covers the UK. UK Biobank’s research ethics committee and Human Tissue Authority research tissue bank approvals mean that researchers wishing to use the resource for approved health research do not need separate ethics approval.

In the EGG+UK Biobank birth weight GWAS[4], multiple births and preterm births were excluded. As gestational age is not available in UK Biobank, an approximation to excluding preterm births was achieved by excluding all births less than 2.2 kg. Estimates for the fetal genetic effect on birth weight at each single nucleotide polymorphism (SNP) was adjusted for the corresponding maternal genetic effect (see below for an explanation of how this was done), in addition to ancestry principal components and centre of recruitment.

As UK Biobank (the largest contributing study to the GWAS) does not have sufficiently powered data on maternal-offspring pairs, the authors of that study[4] derived estimates of the fetal genetic effect on birth weight adjusted for the maternal genotype using a novel method which exploited the reporting by UK Biobank participants of their own birth weight (female and male participants) and the birth weight of their first born offspring (female participants only)[15]. We refer to this as the weighted linear model (WLM) and is described in Box 1. In total, the WLM-GWAS of fetal genotype on birth weight adjusted for maternal genotype used 406,063 participants (101,541 from UK Biobank with own and offspring birth weight, 195,815 from UK Biobank and EGG with own birth weight only and 108,707 from UK Biobank and EGG with maternal genotype and offspring birth weight only; see Figure 1 for more details on the participants)[4]. From the WLM-GWAS, we extracted the estimated fetal per-allele mean difference in birth weight and associated standard error (adjusted for the corresponding maternal effect) for each SNP independently with metabolically favourable adult adiposity (N = 14 SNPs)[10] and adult BMI (N = 76 SNPs)[8] at genome-wide P-value threshold (p<5e-08).

**Figure 1:**
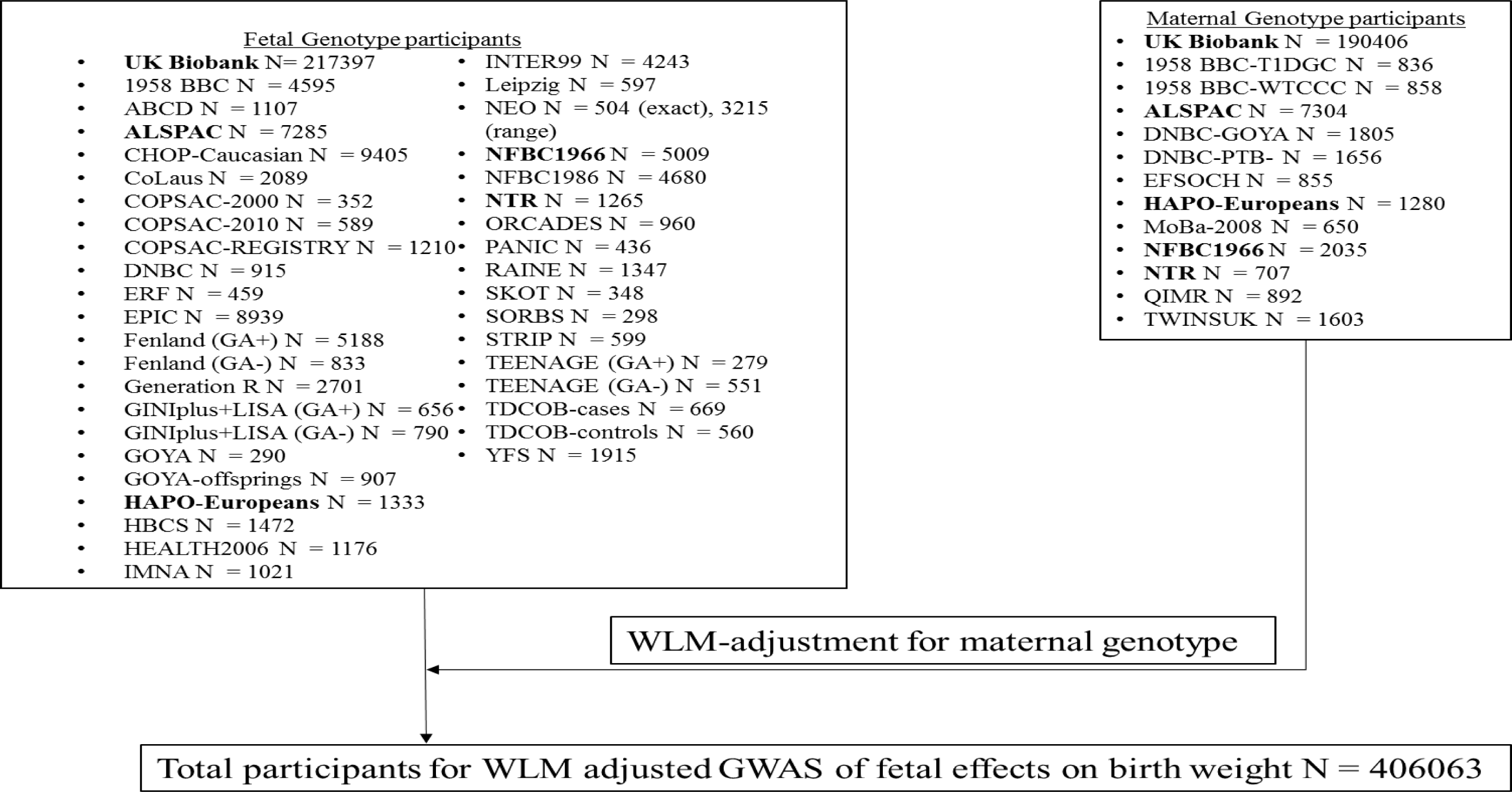
Outline of how all studies in the EGG+UK Biobank meta-analysis contributed to the final GWAS of fetal effects on birth weight[4]. a) Studies in Bold contributed to both the fetal and maternal genotype.

##### Box 1

**Using Structural Equation Modelling (SEM) to estimate maternal and fetal genetic effects on birth weight**

SEM can be used to estimate the fetal genetic effect on own birth weight conditional on maternal genotype in the absence of data from genotyped mother-child pairs[15]. The model uses the participant’s own genotype, own birth weight and their offspring’s birth weight as observed data. For each individual, these observed variables are modelled as functions of two latent (unobserved) variables, the individual’s mother’s genotype and the individual’s offspring’s genotype, which are both correlated 0.5 with the participant’s own genotype. A full description of the structural equation model can be found in Warrington et al 2018[15]. In brief, the model uses the variances and co-variances between the observed variables (own birth weight, offspring birth weight and own genotype), to estimate the parameters of interest including fetal and maternal genetic effects on birth weight. The model is flexible in that it can incorporate a subset of participants with only their own birth weight and own genotype as well as a subset of participant’s with only their own genotype and offspring birth weight. However, fitting the model using full information maximum likelihood is computationally intensive, making it difficult to use at a genome-wide level. Consequently, the authors developed a linear approximation of the full SEM that yielded similar effect estimates and standard errors but was more computationally efficient. This linear approximation, referred to as the weighted linear model (WLM) approximation, combined unadjusted fetal and maternal genetic effect estimates at a single locus, using the following formula

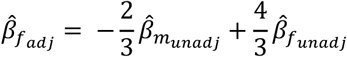

where 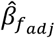 is the estimated fetal genetic effect (adjusted for maternal genotype)on the outcome (in this case birth weight), 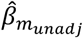 is the unadjusted maternal genetic effect from an unconditional GWAS of maternal genotype and offspring birth weight and 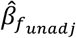 is the unadjusted fetal genetic effect from an unconditional GWAS of fetal genotype and own birth weight [4,15].

Standard errors for the fetal genetic effect (adjusted for maternal genotype) can be calculated, using the following formula.

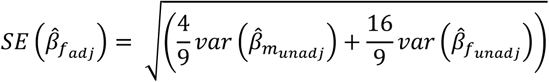

The estimated fetal genetic effect from the WLM,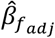, has been shown to be asymptotically equivalent to the estimated effect from a conditional linear model in mother-child pairs, where own birth weight is regressed on own genotype and maternal genotype[4].

#### The four birth cohorts used for analyses of additional birth anthropometric measures and cord-blood outcomes

The Avon Longitudinal Study of Parents and Children (ALSPAC) is a birth cohort that recruited 14,541 pregnancies to women who were resident in and around the city of Bristol in the South West of the UK and who had expected dates of delivery between the 1st of April 1991 to the 31st of December 1992[16,17]. Of these 14,541 there were known live birth outcomes in 13,867 pregnancies to 13,761 women. Please note that the study website contains details of all the data that is available through a fully searchable data dictionary and variable search tool[18]. We used principal components from Hapmap II (release 22) to separate out white Europeans in the genotyped individuals. Maternal genomic data was obtained from the Illumina Human610 Quad Array and fetal data was obtained from the Illumina HumanHap550 Quad Array. The genotypes were imputed using the Haplotype Reference Consortium HRC v1.1 reference panel after quality control (MAF >1%, HWE>1×10^−7^, sex mismatch and kinship errors). In this study we used a maximum of 4,862 unrelated and genotyped mother-child pairs with phenotype data. This cohort contributed to the analyses with the following outcomes: birth weight (included also in the EGG consortium GWAS of birth weight), birth length, birth ponderal index and birth head circumference. Mothers provided written informed consent and ethical approval for the study was obtained from the ALSPAC Ethics and Law Committee and the Local Research Ethics Committees.

Born in Bradford (BiB) is a population-based prospective pregnancy cohort that collected detailed information from 12,450 women who experienced 13,773 pregnancies[19]. The cohort is broadly representative of the obstetric population in Bradford, a city in the North of England, in which approximately half of the births are to mothers of South Asian origin. To be eligible for BiB, women had to have an expected delivery date between March 2007 and December 2010 in the maternity department of the Bradford Royal Infirmary. Participants were recruited primarily at their oral glucose tolerance test (OGTT) appointment, mostly between 26-28 weeks. BiB is a multi-ethnic cohort of mostly White Europeans and South Asians. Ethnicity was based on self-report for most participants, and where that self-report was unavailable, ethnicity reported in GP records was used. For a small number of participants where neither self-report or GP records of ethnicity, South Asian ethnicity was defined using Nam Pechan[19], a computer program for identifying South Asian names[20], and those not identified as South Asian by Nam Pechan were assumed to be White British and included in this study. Maternal and fetal genomic data was obtained from two separate chips, an Illumina HumanCoreExome array and Illumina Infinium Global Screening array (GSA), and genotype data were imputed against HRC r1.1 using Minimac4, after quality control (MAF >1% and HWE>1×10^−6^). In this study we used a maximum of 1,947 unrelated and white European genotyped mother-child pairs with phenotype data. This cohort contributed to the analyses with the following outcomes: birth weight, birth head circumference, birth triceps skinfold thickness, birth subscapular skinfold thickness, sum of skinfold thickness, maternal fasting glucose, maternal 2 hour post-prandial glucose levels, cord-blood insulin, cord-blood leptin (a marker of fetal fat mass[21]) and cord-blood adiponectin. Cord blood was extracted from a vein or artery by the attendant mid-wife at delivery. Samples were refrigerated at 4°C in EDTA tubes until collected by laboratory staff within 12 hours. Samples were then spun, frozen and stored at −80°C. They were transferred to the Biochemistry Department of Glasgow Royal Infirmary for analyses (with no previous thawing), where leptin and adiponectin were measured by a highly sensitive in house ELISA with better sensitivity at lower levels than commercial assays. Insulin was measured using an ultrasensitive solid-phase two-site immunoassay ELISA (Mercodia, Uppsala, Sweden) that does not cross-react with pro-insulin[22]. Laboratory staff were blinded to the participants ethnicity and other characteristics. Ethics approval was obtained for the main platform study and all of the individual sub-studies from the Bradford Research Ethics Committee[19].

The Exeter Family Study of Childhood Health (EFSOCH) is a birth cohort that recruited 1,017 families who were resident in the postcode-defined area of central Exeter between 2000 and 2004, at the Royal Devon and Exeter Hospital[23] from which a total of 993 live births were included in the analyses of this paper. Maternal and fetal genomic data was obtained from the Illumina Infinium HumanCoreExome-24, and the genotypes were imputed against Haplotype Reference Consortium HRC v1.1 reference panel after quality control (MAF >1%, HWE>1×10^−6^, sex mismatch, kinship errors and 4.56 SD from the cluster mean of any sub-populations cluster). In this study we used a maximum of 674 unrelated and genotyped mother-child pairs with phenotype data. This cohort contributed to the analyses with the following outcomes: birth weight (included also in the EGG consortium GWAS of birth weight), birth length, birth ponderal index, birth head circumference, birth triceps skinfold thickness, birth subscapular skinfold thickness, sum of skinfold thickness, maternal fasting glucose and cord-blood insulin. Cord blood was extracted from a vein or artery by the attendant mid-wife at delivery. The blood was stored at 4°C until being collected by the researchers. The cord blood was spun to separate out the plasma which was then stored at −80°C. The plasma was then tested for insulin levels when appropriate at the Regional Endocrine Laboratories (Birmingham, UK) using immunochemiluminometric assays (Molecular Light Technology, Cardiff, U.K.)[23,24]. All mothers and fathers gave informed consent and ethical approval was obtained from the local review committee.

The Hyperglycemia and Adverse Pregnancy Outcomes (HAPO) cohort recruited 28,562 pregnant women between the 1st of July 2000 and the 30th of April 2006 from 15 clinical study centres in 10 countries (United States, Canada, Barbados, United Kingdom, the Netherlands, Thailand, Israel, Australia, Hong Kong and Singapore), four of the centres being in the United States, for their oral glucose tolerance test (OGTT) between 24 and 32 weeks[25]. In total, 25,505 pregnant women underwent OGTT, however only 23,316 women were blind tested (participants were un-blinded if they showed signs of having diabetes i.e. fasting plasma glucose > 5.8 mmol/l or 2 hour glucose > 11.1 mmol/l). The protocol was approved by the institutional review board at each field centre. HAPO is a multi-ethnic cohort, and ethnicity was self-reported by the participants[25]. Maternal and fetal genomic data was obtained from Illumina genome-wide arrays at the Broad Institute (Cambridge, MA) or Johns Hopkins Center for Inherited Disease Research (Baltimore, MD). The genotypes were imputed using SHAPEIT v.2 and IMPUTE2 v.2.3.0 with 1000 Genomes Phase 3 data after quality control as previously described. In this study, we used a maximum of 1,867 unrelated and genotyped mother-child pairs with phenotype data. This cohort contributed to the analyses with the following outcomes: birth weight (included also in the EGG consortium GWAS of birth weight), birth length, birth ponderal index, birth head circumference, birth triceps skinfold thickness, birth subscapular skinfold thickness, sum of skinfold thickness, maternal fasting glucose, maternal 2 hour post-prandial glucose levels and cord-blood c-peptide. Cord blood plasma was extracted at each centre, was stored at −20°C and sent to the Central Laboratory for analysis. A subset of plasma was then stored at −70°C, before being tested for c-peptide using a solid-phase, two-site fluoro-immunometric assay (Autodelfia, Perkin-Elmer, Waltham, Massachusetts, United States). C-peptide has an advantage over insulin in that it is less likely to be destroyed by haemolysis, thus allowing for a more accurate representation of cord insulin levels if haemolysis has occurred in a substantial number of samples[26]. All participants gave written informed consent. An external data and safety monitoring committee provided oversight.

### Data analyses

#### Selecting genetic variants for analyses

The GWAS of metabolically favourable adult adiposity (N=442,278) analysed a composite phenotype characterized by increased body fat percentage and a metabolic profile related to a lower risk of type 2 diabetes, hypertension and heart disease, identified using multivariate GWAS[27] (see Box 2 for more details)[10]. Associations at a total of 14 loci were identified (at p < 5 x 10^−8^), each marked by a SNP at which one allele was associated with higher adult body fat percentage and a “favourable” metabolic profile[10]. We selected these 14 SNPs for our analyses. As these genetic variants were discovered using UK Biobank, and the birth weight GWAS used UK Biobank, there is sample overlap and a potential risk of overfitting and biasing the result towards a confounded association[28]; hence for each SNP, we extracted the effect estimate from the most recent non-UK Biobank GWAS of body fat percentage[29].

##### Box 2

**Deriving the metabolically favourable adiposity phenotype and identifying genetic variants related to this phenotype**

The metabolically favourable adult adiposity genetic variants were identified in a previous study[10] in three steps. In step 1 a GWAS for body fat percentage, as measured by bioimpedance, was performed in the UK Biobank (N = 442,278)[10]. In step 2, a multivariate GWAS was performed combing several metabolic biomarkers together in the same multivariate analyses (i.e. body fat percentage, high density lipoprotein (HDL) cholesterol, adiponectin, sex-hormone binding globulin, triglycerides, fasting insulin and alanine transferase). In order to perform a multivariate GWAS, canonical correlation analyses were conducted as implemented by the metaCCA package in R[27]. Standard univariate GWAS analyses of quantitative traits use linear regression to estimate the linear relationship between one SNP at a time and a single outcome of interest (sometimes adjusting for covariates in the multivariable regression model). Canonical correlation analyses on the other hand involves estimating the maximum correlation between an optimally weighted linear combination of exposure variables and an optimally weighted linear combination of outcome variables. When used in the context of GWAS, this allows one to see how a given genetic variant associates with a linear combination of observed traits, in this particular instance, a combination of traits which index a metabolically favourable vs unfavourable profile. Traditionally, in order to perform multivariate GWAS using canonical correlation analyses, individual level participant data would be needed. However, the metaCCA program allows one to perform canonical correlation analyses using summary results data[27]. In step 3, SNPs that were associated at p < 5 x 10^−8^ with both higher body fat percentage (step 1) and with a metabolically favourable profile in the multivariate GWAS of metabolic traits (step 2) were selected. This was achieved using hierarchical clustering using the pvclust R package, a method which groups of genetic variants are clustered based on their differing associations with observed phenotypes. Finally, the genetic variants identified were then replicated in five obesity cohorts[10].

For BMI, 76 SNPs were identified from the most recent European ancestry GWAS completed prior to the inclusion of the UK Biobank (N=322,154) [8]. Although 77 SNPs were reported in the GWAS[8], one SNP, rs7903146, in *TCF7L2*, is robustly associated with type 2 diabetes[30], and its association with BMI is in part likely due to collider bias, hence it was excluded.

#### Estimating effects of individual genetic variants on birth anthropometric and cord blood markers in the four birth cohorts

Within each of these cohorts, we analysed individual level data using linear regression of birth weight or other perinatal traits on the SNPs (adjusting for gestational age, the child’s sex and maternal genotype). Linear regression analyses were performed separately in each cohort and the results pooled using fixed-effects meta-analysis, resulting in summary data on the association between each SNP and each phenotype. Further information on these cohorts and their contribution to the study can be found in Table 1 and details on how the anthropometric outcomes were measured can be found in Table 2.

**Table 1:** Characteristics of the studies used for analyses of all birth outcomes

|  | ALSPAC | BiB | EFSOCH | HAPO 1<sup>a</sup> | HAPO 2<sup>a</sup> |
| --- | --- | --- | --- | --- | --- |
| Number of Mother-Offspring pairs | 7411 | 3308 | 1022 | 1052 | 815 |
| Country | United Kingdom | United Kingdom | United Kingdom | United States | United States |
| Offspring years of birth | 1991-1993 | 2007-2011 | 2000-2004 | 2001-2006 | 2000-2006 |
| Maternal Age at birth of child (years) | 28.5 (4.8) | 27.1 (6) | 30.4 (5.3) | 32.1 (5.1) | 29.9 (5.4) |
| Maternal pre-pregnancy BMI (kg/m<sup>2</sup>) | 22.9 (3.8) | 26.6 (5.9) | 24 (4.4) | 24.2 (4.6) | 24.6 (5.3) |
| Gestational age at delivery (weeks) | 39.6 (1.7) | 39.7 (1.8) | 39.9 (1.5) | 40 (1.2) | 40 (1.2) |
| Offspring sex (% male) | 49.8 | 51.6 | 51.6 | 47.9 | 50.9 |
| Mothers smoking (%) | 17.2 | 33.1 | 13.3 | 12.9 | 15.1 |
| Birth Weight (g) | 3495 (471) | 3439 (482) | 3513 (476) | 3543 (509) | 3540 (431) |
| Birth Length (cm) | 50.9 (2.2) | NA | 50.3 (2.1) | 50.5 (2.2) | 51.8 (2.5) |
| Birth Ponderal Index (kg/m<sup>3</sup>) | 26.4 (2.7) | NA | 27.7 (2.6) | 27.4 (3.3) | 25.4 (3.3) |
| Birth Head Circumference (cm) | 35 (1.4) | 34.7 (1.4) | 35.2 (1.3) | 34.9 (1.6) | 34.9 (1.4) |
| Birth Triceps Skinfolds (mm) | NA | 5.2 (1.1) | 4.9 (1.1) | 4.1 (0.8) | 4.1 (0.9) |
| Birth Subscapular Skinfolds (mm) | NA | 4.9 (1.1) | 4.9 (1.2) | 4.6 (1) | 4.3 (1) |
| Sum of Birth Skinfolds (mm) | NA | 10.1 (2.1) | 9.7 (2.1) | 13.1 (2.5) | 12.3 (2.4) |
| Cord-blood C-Peptide (µg/mL)<sup>b</sup> | NA | NA | NA | 0.9 (0.7-1.3) | 0.9 (0.7-1.3) |
| Cord-blood insulin (pg/mL)<sup>b</sup> | NA | 3.5 (2.1-5.8) | 37.6 (26-60) | NA | NA |
| Cord-blood leptin (ng/mL)<sup>b</sup> | NA | 7.3 (4-13.1) | NA | NA | NA |
| Cord-blood adiponectin (µg/ml)<sup>b</sup> | NA | 33.3 (26.3-42.7) | NA | NA | NA |
| Fasting Glucose (mmol/l) | NA | 4.4 (0.42) | 4.35 (0.38) | 4.58 (0.37) | 4.51 (0.34) |
| 2 hour postload glucose (mmol/l) | NA | 5.43 (1.3) | NA | 6.02 (1.2) | 6.06 (1.19) |

- a) For HAPO 1, genetic data was stored and analysed at the University Feinberg School of Medicine, Chicago. For HAPO 2, genetic data was stored and analysed at the University of Exeter. These were non-overlapping samples of European mothers and babies. - b) For the cord-blood outcomes, because they have a non-standard distribution, the median and interquartile range are displayed. For all other outcomes, the mean value and the standard deviation are displayed.

**Table 2:** Measurement of birth anthropometric traits in selected cohorts

| Trait | ALSPAC[16,17] | BiB[19,43] | EFSOCH[23] | HAPO[44] |
| --- | --- | --- | --- | --- |
| Birth Weight | Extracted from clinical records | Extracted from clinical records | Calibrated electronic scale | Calibrated electronic scale |
| Birth Length | Extracted from clinical records | NA | Standardized plastic board | Standardized plastic board |
| Ponderal Index | Calculated | NA | Calculated | Calculated |
| Head Circumference | Extracted from clinical records | Standardized measuring tape | Standardized measuring tape | Standardized measuring tape |
| Triceps Skinfold | NA | Calipers | Calipers | Calipers |
| Subscapular Skinfold | NA | Calipers | Calipers | Calipers |
| Sum of Skinfolts | NA | Calculated | Calculated | Calculated |

Though our analyses of birth weight and other perinatal traits was restricted to White European ancestry participants, the cohorts used came from different locations and used different methods to measure perinatal traits (especially cord-blood traits). Therefore, to test for between study heterogeneity, we performed Cochran’s Q test and estimated I^2^ for each meta-analyses[31].

#### Combining effects of individual SNPs on birth weight and other outcomes using summary data

To maximise power, we used inverse variance weighted meta-analysis to combine the effects of multiple SNPs (i.e. 14 metabolically favourable adult adiposity SNPs or 76 adult BMI SNPs) on birth weight (or other perinatal traits). This procedure reproduces estimates of the effect of metabolically favourable adult adiposity and BMI allele scores on birth outcomes without requiring individual level data. We did this by meta-analysing the fetal SNP-birth weight (or other perinatal trait) associations and weighting by the adult adiposity or BMI effects[32]. We decided to weight the SNPs as we assumed that any SNP with a large effect on metabolically favourable adult adiposity or BMI would have a correspondingly large effect on birth weight if it influences fetal growth.

Specifically, we multiplied each of the fetal effects on birth weight (or other perinatal traits) for the 14 metabolically favourable adult adiposity SNPs by their per allele association coefficient with percent fat mass[29] (see selecting genetic variants above for the source of this information). Similarly, we multiplied each of the fetal effects on birth weight (or other perintal traits) for the 76 adult BMI by their per allele association coefficient with BMI[8]. For the main analyses, we used the EGG + UK Biobank summary results for the fetal effect on birth weight adjusted for maternal genotype (see Box 1 above). For simplicity, hereafter we refer to these pooled genetic effects as “pooled genetic scores”.

To enable comparison between the two pooled genetic scores, we scaled their effects on birth weight and the other perinatal traits to be equal to 1 SD of the score. To obtain the SD, we generated genetic scores for metabolically favourable adult adiposity and BMI using individual level data in 479,434 UK Biobank participants. For each participant, the adiposity raising alleles for each of the 14 metabolically favourable adult adiposity SNPs were each weighted by their effect on body fat percentage and added together, divided by the sum of the weights and multiplied by the number of SNPs (a similar method was used for the 76 adult BMI SNPs). We then examined the distribution of the score across the participants. The SD of the metabolically favourable adult adiposity score was 2.6 weighted alleles, the SD of the adult BMI score was 5.5 weighted alleles. These SD values were then used to scale the pooled genetic score associations with birth weight, which we generated from summary data.

Birth weight and all other perinatal traits were standardized within each cohort separately when estimating the metabolically favourable adiposity and BMI pooled genetic score effects, in order to make the estimates for each trait comparable with each other. For the EGG+UK Biobank estimates, we converted the results back to grams by multiplying by 484 (the average SD for birth weight in grams for 18 studies in an early birth weight GWAS[33]).

#### Additional analyses

We explored the plausibility of the *a priori* assumption that metabolically favourable adult adiposity and BMI alleles have effects on birth weight proportional to their effects on adult adiposity. To do this, we plotted scatter plots of the fetal effect estimate of each of the SNPs on birth weight (y-axis)[4] and on adult adiposity traits (either body fat percentage or BMI; x-axis)[8,29]. We used inverse variance weighting (i.e. multiplying the SNP-adult trait estimate by the inverse of the variance for the SNP-birth weight effect estimates) when fitting regression lines to each plot[34]. This approach also allowed us to empirically test for outliers from any relationship that may indicate pleiotropy.

## Results

### Fetal genetic variants that predispose to higher metabolically favourable adiposity are more strongly associated with higher birth weight than those that predispose to higher BMI

Children inheriting more metabolically favourable adiposity alleles were heavier at birth (10 grams (95% CI: 7 to 13) difference in mean birth weight per 1 SD increase in the pooled genetic score; Figure 2). In contrast, children inheriting more BMI alleles were heavier at birth but the effect was smaller (4 grams (95% CI: 0 to 8) difference in mean birth weight per 1 SD increase in the weighted allele score; Figure 2). These effects were independent of the maternal genotype.

**Figure 2:**
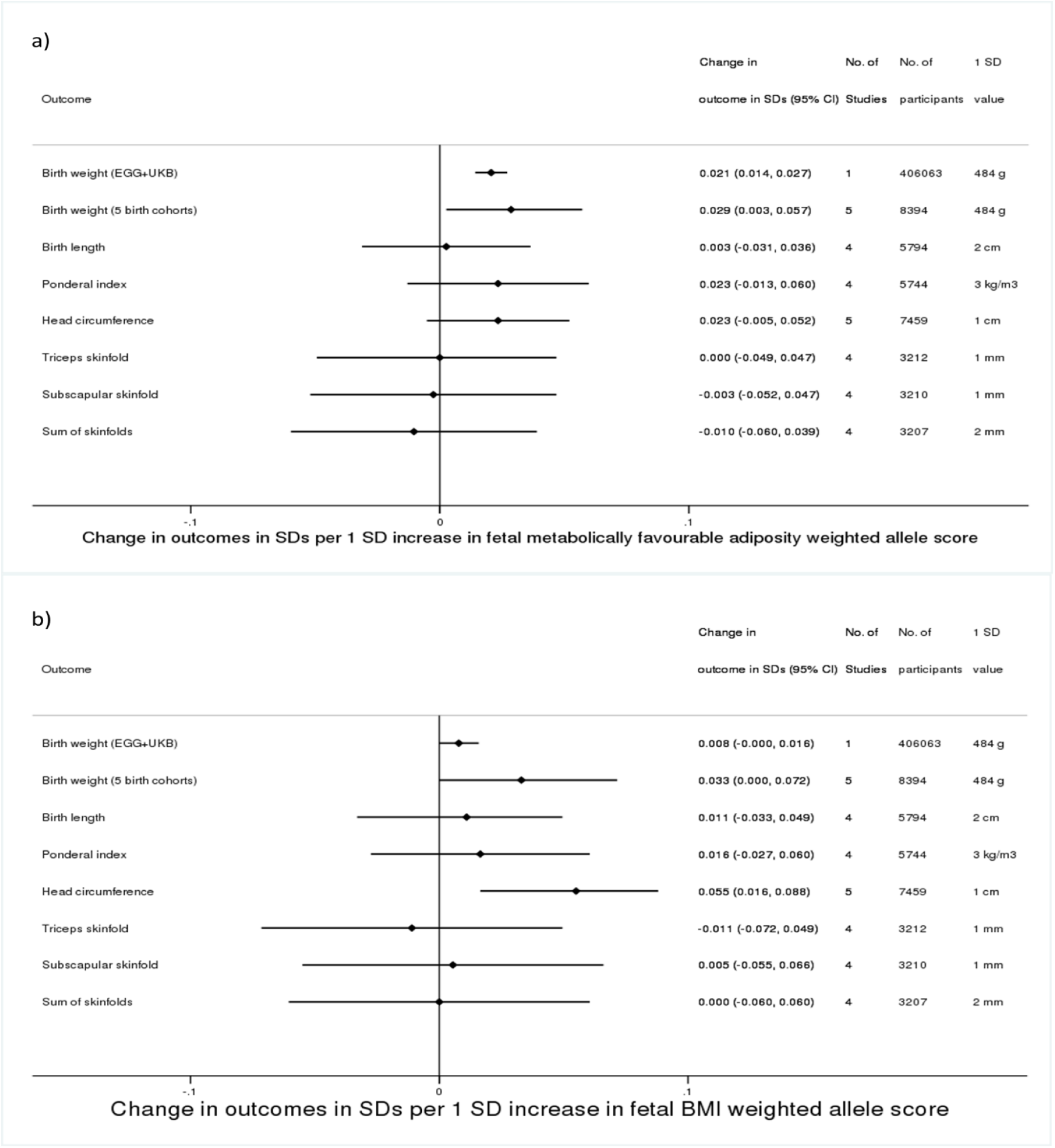
Genetic associations of fetal a) metabolically favourable adiposity and b) BMI pooled genetic scores with birth anthropometric outcomes. a) For Birth weight and Head circumference 5 studies are mentioned; this equates to ALSPAC, BiB, EFSOCH, HAPO 1 and HAPO 2 b) For Birth weight (EGG+UKB), the number of participants is the number involved in the GWAS of own birth weight adjusted for maternal genotype using the WLM (that is 101,541 UKB participants who reported their own birth weight and birth weight of their first child, 195,815 UKB and EGG participants with own birth weight data, and 108,707 UKB and EGG participants with offspring birth weight data)[4]

The effects of both fetal metabolically favourable adiposity and BMI pooled genetic scores on birth weight and other perinatal anthropometric outcomes in the four mother-child pair cohorts were consistent in terms of magnitude and direction with the results from the main EGG+UK Biobank results, though with wider confidence intervals due to lower sample sizes (Figure 2).

All of the fetal metabolically favourable adiposity or BMI pooled genetic score estimates for the cord-blood markers had wide confidence intervals owing to the small sample sizes, making it difficult to say if there was any effect (Figure 3). There was substantial between study heterogeneity for the fetal pooled genetic score related to higher metabolically favourable adiposity effect on cord-insulin levels (Cochrane’s Q = 5.92 (d.f. = 1), I^2^=83%), though there was minimal between study heterogeneity for the other outcomes.

**Figure 3:**
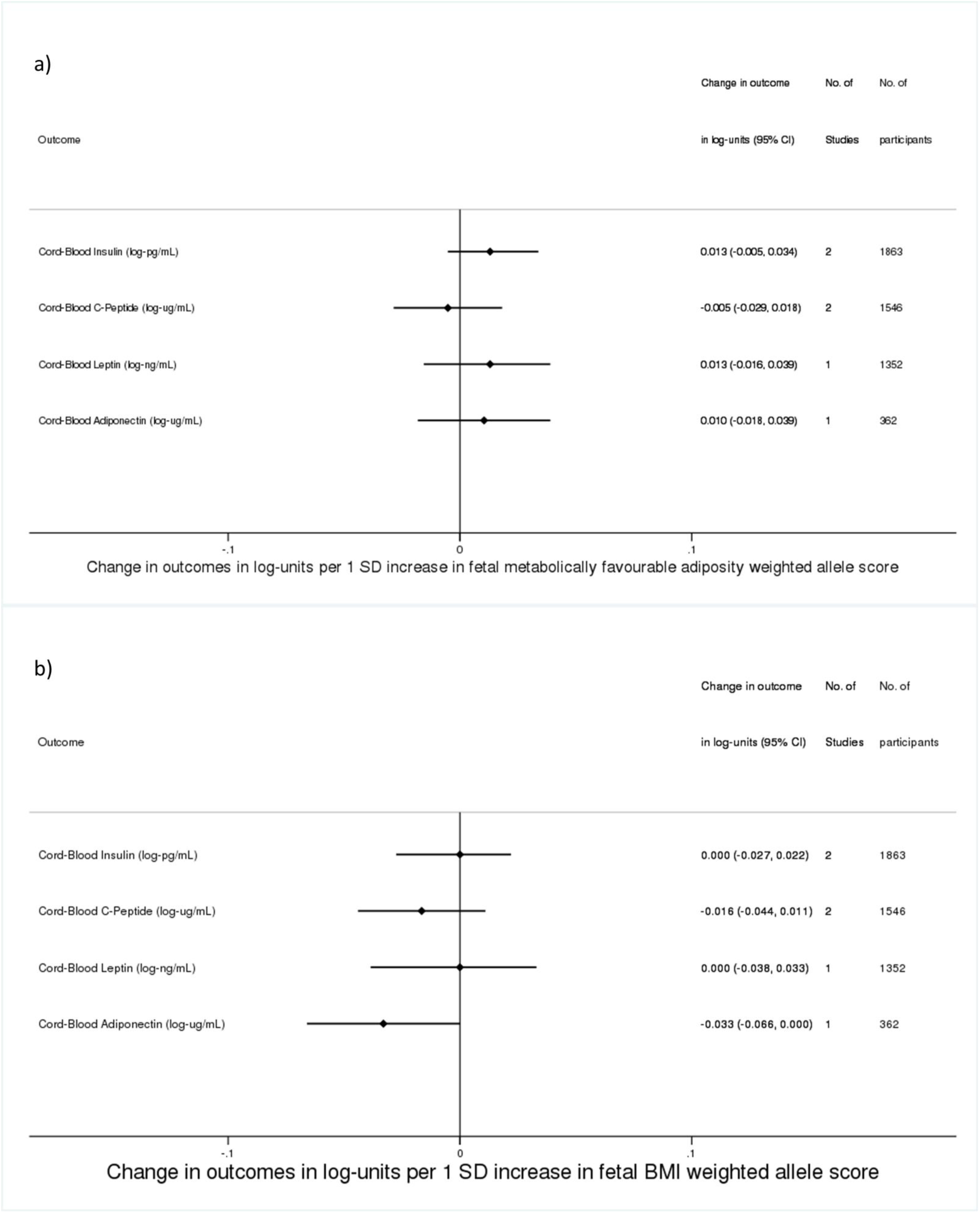
Genetic associations of fetal a) metabolically favourable adiposity and b) BMI pooled genetic scores with cord-blood outcomes

### The effect of fetal genetic variants that predispose to higher metabolically favourable adiposity on birth weight is proportional to their effect on adult adiposity

We plotted the SNP-birth weight effects against both SNP-body fat percentage effects (Figure 4) and SNP-BMI effects (Figure 5). The metabolically favourable adiposity alleles were associated with birth weight and adult measures of adiposity in a dose-dependent manner, with the alleles with the largest effects on birth weight having the largest effect on body fat percentage (R^2^ = 0.29). In contrast, there was no clear evidence that the BMI alleles were associated with birth weight and any adult measures of adiposity in a dose-dependent association (R^2^ = −0.01), suggesting the presence of substantial across SNP heterogeneity in the BMI alleles effects on birth weight and adult measures of adiposity.

**Figure 4:**
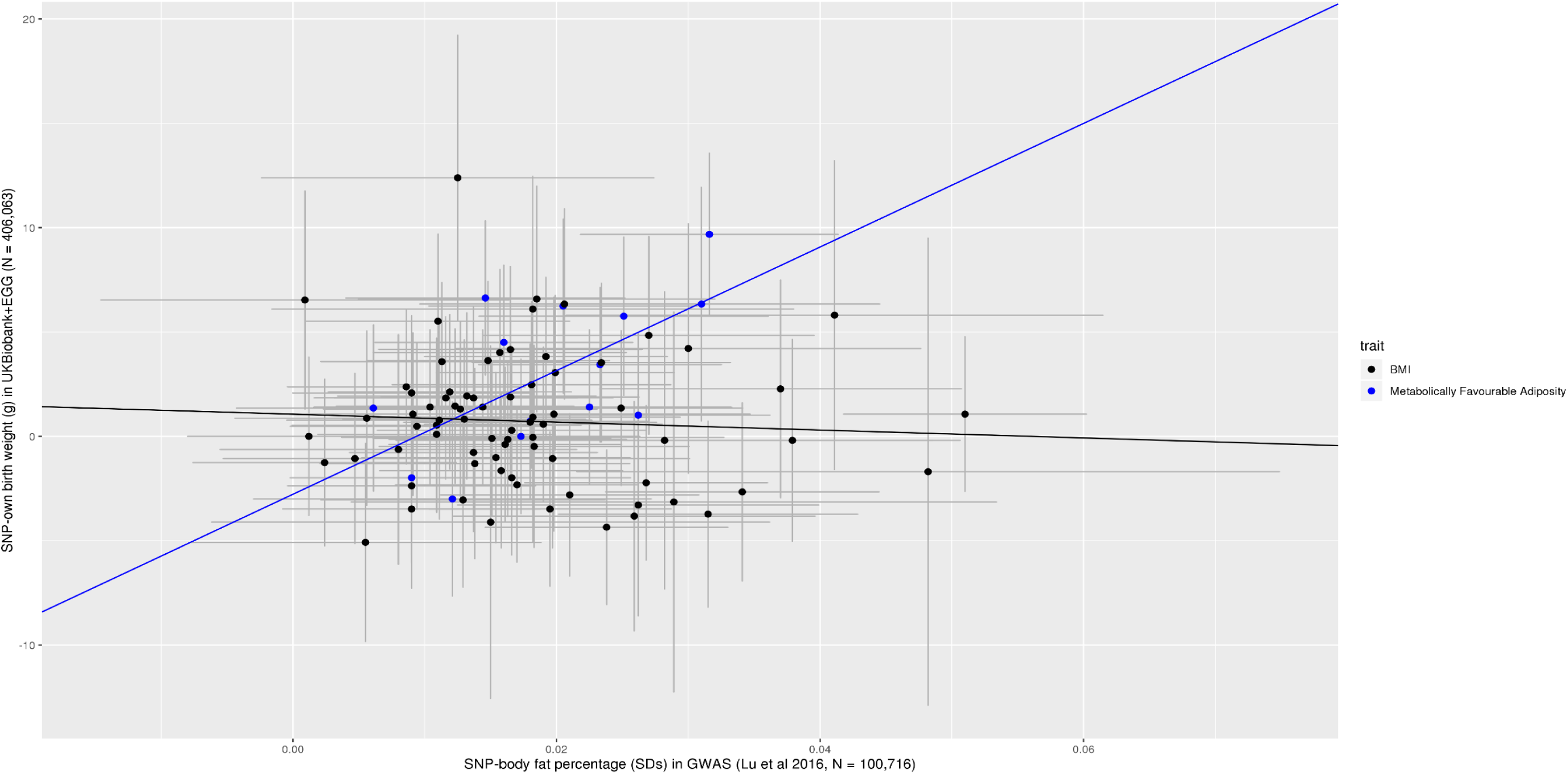
Scatter plot of SNP effects on body fat percentage (x-axis) and SNP effects on birth weight (y-axis) a) Inverse-variance weighting (multiplying the by the inverse of the standard error for the SNP-birth weight per allele association) was used when fitting regression lines to each set of SNPs b) The error bars represent the 95% confidence intervals

**Figure 5:**
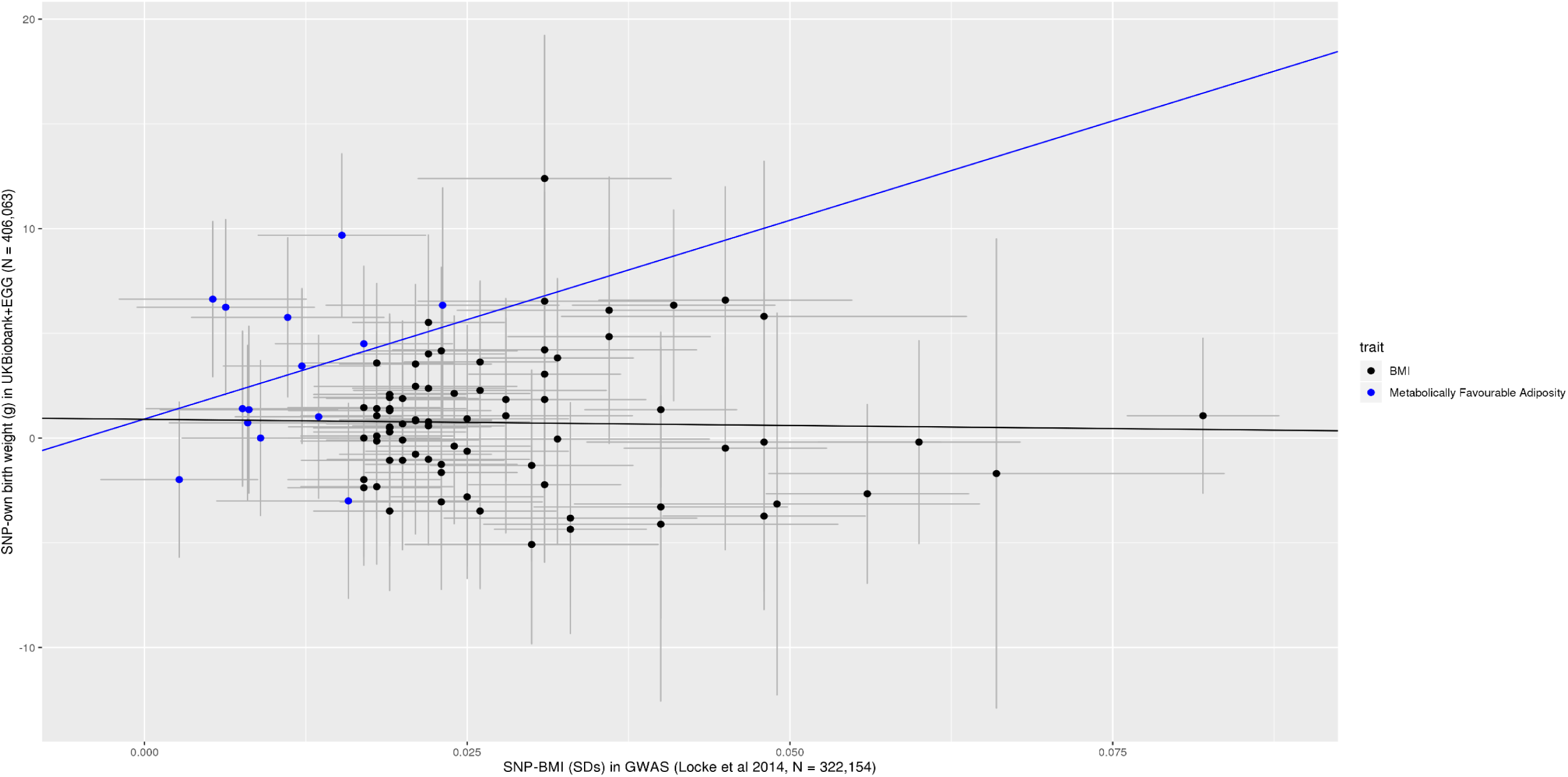
Scatter plot of SNP effects on BMI (x-axis) and SNP effects on birth weight (y-axis) a) Inverse-variance weighting (multiplying the by the inverse of the standard error for the SNP-birth weight per allele association) was used when fitting regression lines to each set of SNPs b) The error bars represent the 95% confidence intervals

## Discussion

Using data on more than 400,000 individuals, we have shown that a higher fetal genetic score for metabolically favourable adiposity in adulthood is associated with higher birth weight. A higher fetal genetic score for adult BMI is also associated with higher birth weight, but with a smaller effect size. The metabolically favourable adiposity SNPs with larger adult adiposity effects tended to show larger effects on birth weight (Figure 4). In contrast, there was no correlation for the BMI SNPs between their adult adiposity effect sizes and their birth weight effect sizes (Figure 5), suggesting substantial heterogeneity of effects of fetal BMI genetic variants on fetal growth. The effect of the fetal genetic variants on other birth anthropometric outcomes was directionally consistent with their effects on birth weight. For cord-blood markers, effects of both metabolically favourable adiposity and BMI genetic variants were close to the null. However, we had limited power for results beyond birth weight and so these results should be treated with caution until replicated in larger studies.

This study provides further evidence of the importance of fetal genetics in determining offspring birth weight. Early studies using BMI genetic variants, without adjusting for the maternal genotype, found little evidence of an effect on birth weight[5]. This was corroborated by a GWAS of BMI during the early stages of childhood; the variants identified to be associated with BMI in early life had minimal effect on birth weight[35]. However, a recent large scale GWAS, using LD score regression with a previous GWAS, found a small genetic correlation between BMI and fetal genetic effect on own birth weight, adjusted for maternal genotype (r_g_ = 0.12)[4]. Our scatter plot analysis showed evidence of a dose response effect of metabolically favourable adiposity associated SNPs on birth weight, but this was not seen for BMI associated SNPs. These findings are consistent with the observation that most SNPs identified in the BMI GWAS map to genes expressed in the brain[8], in particular regions of the brain like the insula and substatia nigra that are implicated in reward and addiction processes[36]. Therefore these genetic variants might influence adult adiposity by influencing a neuronal pathway which in turn regulates postnatal appetite rather than general body growth.

This study also provides further evidence of the interplay between maternal and fetal genetic effects on fetal growth. In a previous Mendelian Randomisation study we have shown that higher maternal (genetically instrumented) metabolically favourable adiposity leads to lower offspring birth weight[13]. Considering those findings with the ones from the current paper our work suggests that mothers with metabolically favourable adiposity alleles will on average have more metabolically favourable adiposity and fetal intrauterine exposure to this would result in them having a lower birth weight. At the same time, the fetus of these women will have more metabolically favourable adiposity alleles which will result in them having higher birth weight. Completely separating the maternal from fetal effects may be difficult but highlights the importance for adjusting for maternal genetic variants as we have done here, and for the fetal genetic variants in Mendelian Randomisation studies of maternal pregnancy exposures as we have done here. A recent GWAS of fetal genetic effects on own birth weight, adjusted for maternal genotype, found that genetic predisposition to higher adult fasting glucose levels was associated with lower birth weight, possibly due to decreased capacity for insulin secretion[4]. However, metabolically favourable adiposity variants are linked to greater insulin sensitivity rather than greater insulin secretion[10]. As insulin has been shown to act as a growth factor *in utero*[37], a possible mechanism for the effect of fetal metabolically favourable adiposity alleles with higher birth weight may be greater insulin sensitivity, allowing for a greater growth response to insulin secretion. It is also possible that fetal metabolically favourable adiposity alleles allow for greater fetal fat mass accumulation and hence greater birth weight. This possibility is supported by the fact that several of the metabolically favourable adiposity loci, in particular *PPARG*, have previously been found to be associated with adipocyte differentiation[38]. Our analyses of cord-blood insulin and neonatal anthropometric traits aimed to assess evidence that higher fetal insulin secretion and/or fat mass might underlie the birth weight effects, but further studies with larger samples are needed to provide sufficient statistical power.

### Strengths and Limitations

This study is the first to investigate the fetal effects of the recently discovered metabolically favourable adiposity associated genetic variants on fetal outcomes independent of the corresponding maternal genetic effects. It is also the largest study to investigate the effects of BMI associated genetic variants on fetal outcomes. To do this we attempted to use all available relevant data from multiple independent cohorts, increasing the certainty of our findings.

Despite using all available mother-child cohorts with relevant data, our analyses had relatively low statistical power for outcomes other than birth weight, hence there is a need for larger cohort studies and/or GWAS of other neonatal outcomes.

Also, there was substantial overlap between the samples of the metabolically favourable adiposity and birth weight GWAS’s, which could bias our estimates due to statistical overfitting. To mitigate any impact of this bias, we have weighted our score using estimates from an external independent population.

A further limitation to this study is the low response rate for UK Biobank (5.5%)[39] and the self-report of own birth weight in UK Biobank and some of the other studies included in EGG. A highly selected cohort such as UK Biobank can result in selection bias in genetic analyses[40,41]. However, whilst self-report of birth weight is likely to have some measurement error, it is unlikely to affect the fetal genotype itself. The fact that the results for birth weight in EGG+UK Biobank were consistent with the results for birth weight of the four mother-offspring pair cohorts combined suggests that this bias is unlikely to have materially affected the results. For this study, we limited ourselves to European ancestry individuals, though relevant studies involving African ancestry individuals have been published[42], and additional studies involving non-European populations will be important going forward.

We have described the results of adiposity related genetic variants influencing birth weight, as the genetic variants cannot be influenced by the confounding of many social and environmental factors that confound conventional associations of non-genetic factors with outcomes, nor can genetic variants be influenced by existing disease. We limited analyses to participants of European origin and the GWAS adjusted for principal components and centre of recruitment, which should limit any confounding of the genetic effect due to population stratification.

However, whilst we do not think these findings are confounded, we acknowledge that we do not know much about the function of the genes included in the pooled genetic scores used here and so do not know how they influence adiposity or birth weight.

In conclusion, our results suggest that fetal genetic predisposition to higher metabolically favourable adult adiposity is associated with higher birth weight, and that this effect is stronger than that of genetic predisposition to higher general adult adiposity, the latter proxied by BMI. Larger population samples are needed to investigate the effects on other birth anthropometric outcomes and cord-blood markers, in order to elucidate the mechanisms underlying these effects.

## Abbreviations

ALSPAC: (Avon Longitudinal Study of Parents and Children),
BiB: (Born in Bradford),
EFSOCH: (Exeter Family Study of Childhood Health),
EGG: (Early Growth Genetics consortium),
HAPO: (Hyperglycaemia and Adverse Pregnancy Outcomes),
SEM: (Structural Equation Modelling),
WLM: (Weighted Linear Model)

## Acknowledgements

This research has been conducted using the UK Biobank Resource under application number 7036. We would like to thank the participants and researchers from the UK Biobank who contributed or collected data. We are extremely grateful to all of the families who took part in ALSPAC, the midwives for their help in recruiting them, and the whole ALSPAC team, which includes interviewers, computer and laboratory technicians, clerical workers, research scientists, volunteers, managers, receptionists and nurses. We are grateful to everyone involved in the Born in Bradford study. This includes the families who kindly participated as well as the practitioners and researchers all of whom have made Born in Bradford happen. We are also grateful for the families that took part in EFSOCH and the researchers that collected data. We are grateful to the Genetics of Complex traits team at the University of Exeter, for their assistance in learning the methods and navigating the study data; in particular we are grateful to Francesco Casanova who helped with the data extraction for BiB.

## Data Availability

Our study used both published summary results (i.e. taking results from published research papers and websites) and individual participant cohort data as follows:

The data for the GWAS of BMI is available here. https://portals.broadinstitute.org/collaboration/giant/index.php/GIANT_consortium_data_files

The data for the GWAS of body fat percentage is available here. https://walker05.u.hpc.mssm.edu

The data for the GWAS of birth weight is available here. https://egg-consortium.org/birth-weight-2019.htm

The references to those published data sources are provided in the main paper.

We used individual participant data for the genetic association analyses from the UK Biobank, ALSPAC, BiB, EFSOCH and HAPO cohorts.

The data in UK Biobank, ALSPAC and BiB are fully available, via managed systems, to any researchers. The managed system for both studies is a requirement of the study funders but access is not restricted on the basis of overlap with other applications to use the data or on the basis of peer review of the proposed science.

UK Biobank. Full information on how to access these data can be found here - https://www.ukbiobank.ac.uk/using-the-resource/

ALSPAC. The ALSPAC data management plan (http://www.bristol.ac.uk/alspac/researchers/data-access/documents/alspac-data-management-plan.pdf) describes in detail the policy regarding data sharing, which is through a system of managed open access. The steps below highlight how to apply for access to the data included in this paper and all other ALSPAC data.

1. Please read the ALSPAC access policy (PDF, 627kB) which describes the process of accessing the data and samples in detail, and outlines the costs associated with doing so.
2. You may also find it useful to browse the fully searchable ALSPAC research proposals database, which lists all research projects that have been approved since April 2011.
3. Please submit your research proposal for consideration by the ALSPAC Executive Committee. You will receive a response within 10 working days to advise you whether your proposal has been approved.

If you have any questions about accessing data, please.

BiB. Full information on how to access these data can be found here - https://borninbradford.nhs.uk/research/how-to-access-data/

HAPO. For access to the data used in this study, please contact Dr. Rachel Freathy and Prof. William Lowe Jr. The website describing the study and other data available is https://www.ncbi.nlm.nih.gov/projects/gap/cgi-bin/study.cgi?study_id=phs000096.v4.p1

If you have further questions, please email Dr William Lowe at

EFSOCH. Requests for access to the original EFSOCH dataset should be made in writing in the first instance to the EFSOCH data team via the Exeter Clinical Research Facility.

## Author Contributions

DAL and RMF designed this study, with WDT further developing the design.

DAL (ALSPAC), ATH and BAK (EFSOCH), DS and WLL (HAPO) and GS, RA, DM and DAL (BiB) contributed to the collection of and management of cohort data.

ARW, RNB and JT prepared the imputed genotype and phenotype data for the UK Biobank sample of European ancestry participants.

TMF, HY and YJ performed a multivariate GWAS to discover favourable adiposity genetic variants.

NW and DME performed a Weighted Linear Model GWAS using the UK Biobank + EGG genetic and birth weight data.

WDT, RMF, DAL and M-CB wrote the analysis plan, and WDT undertook most of the analyses with support from JT, M-CB, RB, RMF and DAL.

AK carried out data analyses using the HAPO dataset in Chicago, and in this task he was supervised by DS.

WDT wrote the first draft of the paper with support from M-CB, RMF and DAL; all authors read and made critical revisions to the paper.

WDT, RMF and DAL act as guarantors for the paper’s integrity

